# Learning what is irrelevant or relevant: Expectations facilitate distractor inhibition and target facilitation through distinct neural mechanisms

**DOI:** 10.1101/565069

**Authors:** Dirk van Moorselaar, Heleen A. Slagter

## Abstract

It is well known that attention can facilitate performance by top-down biasing processing of task-relevant information in advance. Recent findings from behavioral studies suggest that distractor inhibition is not under similar direct control, but strongly dependent on expectations derived from previous experience. Yet, how expectations about distracting information influence distractor inhibition at the neural level remains unclear. The current study addressed this outstanding question in three experiments in which search displays with repeating distractor or target locations across trials allowed observers to learn which location to selectively suppress or boost. Behavioral findings demonstrated that both distractor and target location learning resulted in more efficient search, as indexed by faster response times. Crucially, benefits of distractor learning were observed without target location foreknowledge, unaffected by the number of possible target locations, and could not be explained by priming alone. To determine how distractor location expectations facilitated performance, we applied a spatial encoding model to EEG data to reconstruct activity in neural populations tuned to the distractor or target location. Target location learning increased neural tuning to the target location in advance, indicative of preparatory biasing. This sensitivity increased after target presentation. By contrast, distractor expectations did not change preparatory spatial tuning. Instead, distractor expectations reduced distractor-specific processing, as reflected in the disappearance of the Pd ERP component, a neural marker of distractor inhibition, and decreased decoding accuracy. These findings suggest that the brain may no longer process expected distractors as distractors, once it has learned they can safely be ignored.

**Significance statement:** We constantly try hard to ignore conspicuous events that distract us from our current goals. Surprisingly, and in contrast to dominant attention theories, ignoring distracting, but irrelevant events does not seem to be as flexible as is focusing our attention on those same aspects. Instead, distractor suppression appears to strongly rely on learned, context-dependent expectations. Here, we investigated how learning about upcoming distractors changes distractor processing and directly contrasted the underlying neural dynamics to target learning. We show that while target learning enhanced anticipatory sensory tuning, distractor learning only modulated reactive suppressive processing. These results suggest that expected distractors may no longer be considered distractors by the brain once it has learned that they can safely be ignored.

## Introduction

Attention can top-down bias sensory processing of task-relevant information in advance (Battistoni, Stein, & Peelen, 2017). Yet, it is currently debated whether distractor inhibition is under similar direct control. That is, in contrast to the widely accepted view that alpha-band oscillations implement top-down inhibition (Foxe & Snyder, 2011; Jensen & Mazaheri, 2010), recent behavioral findings suggest that distractor inhibition strongly relies on previous experiences with visual distractions (Noonan et al., 2016; Wang & Theeuwes, 2018a). This has led to the proposal that distractor filtering is not resolved through top-down inhibition, but instead through expectation-dependent suppression of distractor processing. In this view, consistent with predictive processing theories, any expected stimulus, whether relevant or irrelevant, is suppressed (explained “away”), unless attention releases it from expectation-dependent suppression (Noonan, Crittenden, Jensen, & Stokes, 2018).

Behavioral studies have established that (implicit) learning where targets or distractors are most probable influences visual selection (Ferrante et al., 2018; Geng & Behrmann, 2002, 2005; Jiang, 2018; Wang & Theeuwes, 2018b). Spatial probability learning arguably generates expectations where stimuli are most likely to occur, which may in turn bias attention. Predictive processing theories postulate that such expectations, which need not necessarily be conscious, attenuate sensory responses (Friston, 2009; Rao, 2005). Yet, ideas differ whether expectations exert their influence already in advance (Fiser et al., 2016) or alternatively, only become apparent after stimulus presentation (Bar et al., 2006; Rao & Ballard, 1999). In line with the former notion, a recent study demonstrated expectation-dependent anticipatory sharpening of stimulus representations (Kok, Mostert, & De Lange, 2017). In addition to expectation, preparatory attention also induces stimulus templates in sensory cortex that facilitate target selection (Myers et al., 2015). Expectation and attention also interact, such that top-down biasing, as reflected in pre-stimulus alpha-lateralization, is most pronounced when targets also most likely occur at the cued, task-relevant location (Alilović, Timmermans, Reteig, van Gaal, & Slagter, 2018). Thus, task relevant expectations may bias corresponding visual regions in advance to facilitate goal-directed behavior. Yet, how distractor-specific expectations help resolve interference - through modulating pre-stimulus activity representing the distracting information and/or post-distractor processing – is unclear.

Notably, a recent study reported no changes in pre-stimulus alpha-lateralization as a function of distractor location learning despite clear behavioral evidence for distractor suppression (Noonan et al., 2016). This absence of any distractor-related alpha-lateralization is surprising in light of the prevailing view that alpha oscillations implement top-down inhibition of activity in irrelevant visual networks (Foxe & Snyder, 2011; Jensen & Mazaheri, 2010). Yet, the vast majority of this work examined inhibition in binary designs, where one of two features was irrelevant. Hence, observed effects could also reflect secondary inhibition related to more attention to task-relevant features, or simply attending away, rather than top-down inhibition per se (Foster & Awh, 2018).

The present study was designed to characterize the neural mechanisms underlying learned distractor suppression and directly contrast these with target learning. Specifically, we aimed to establish whether learned suppression is purely reactive, or already evident in anticipation of expected distractors. For this purpose, in two behavioral and one electroencephalography (EEG) experiment, across visual searches, either the distractor or the target location was repeated, allowing observers to learn which location to selectively ignore or select (see Fig. 1A). As target locations could not be predicted in the distractor-repeat condition, any observed effect of distractor learning is unlikely to be driven by more attention to expected target locations. Changes in preparatory and stimulus-induced activity and neural representation were examined using inverted encoding modeling, ERP, and multivariate pattern analyses. Based on recent studies showing expectation- and attention-dependent anticipatory sharpening of stimulus representations (Foster, Sutterer, Serences, Vogel, & Awh, 2017; Samaha, Sprague, & Postle, 2016), we expected target learning to enhance reconstruction of target locations before search display onset. However, if and how, distractor learning also changed spatial distractor tuning in advance or only modulated post-distractor processing was an open question.

**Figure 1.**
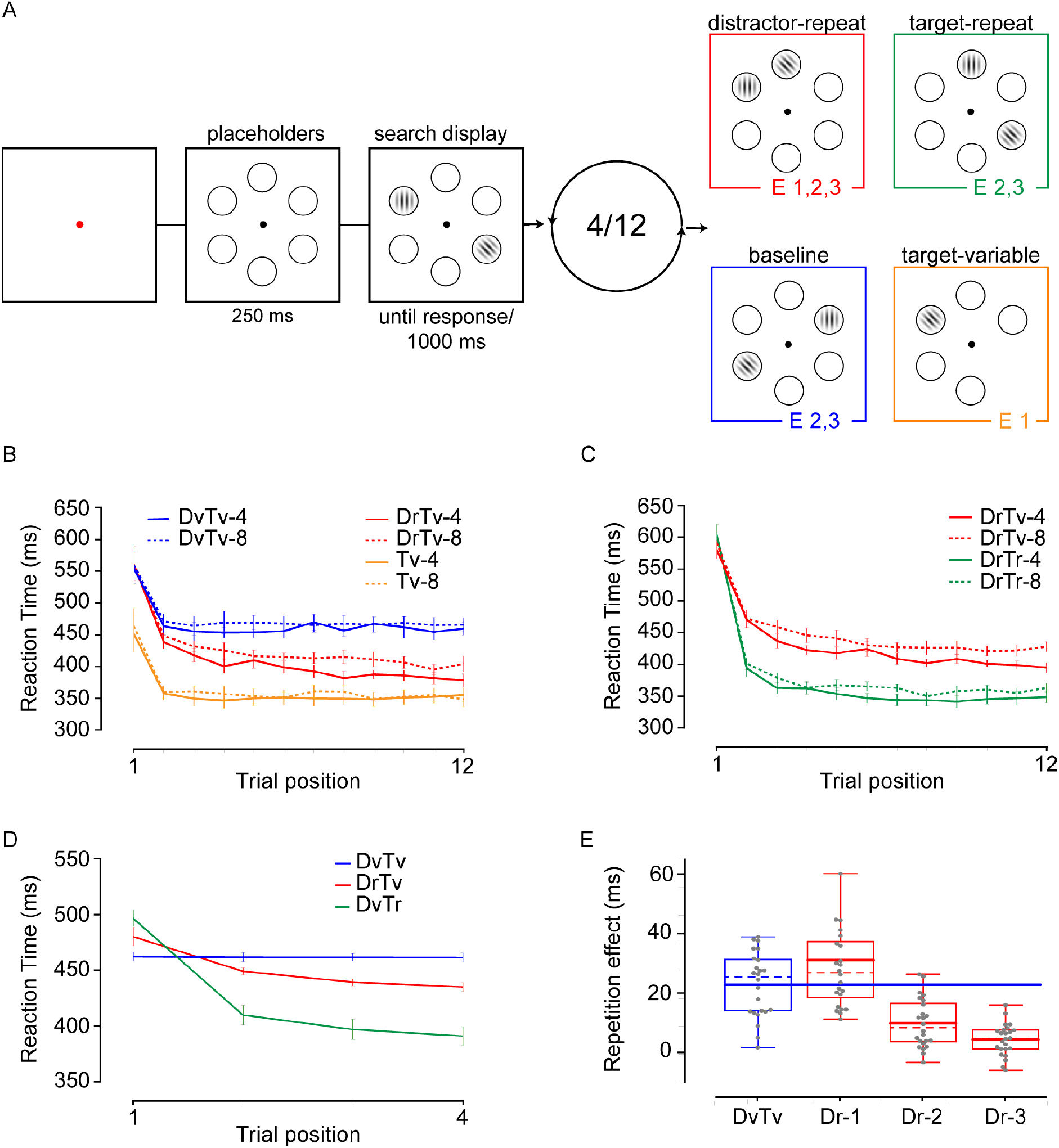
Task design and behavioral findings of Experiments 1-3. (**A**) A trial sequence of Experiment 1-3. In each condition, in each trial of a sequence of trials, participants had to indicate the orientation (left or right) of a target gabor. In all conditions, except the target-only variable (Tv) condition, the target was accompanied by a distractor (a horizontally or vertically oriented gabor). In the target-repeat (DvTr) and distractor-repeat (DrTv) conditions, the location of the target (Tr) or the distractor (Dr) was repeated over trials in a sequence. In the baseline (DvTv) condition, the target and distractor location varied from trial to trial. In the target-only variable condition (Tv), the location of the target also varied from trial to trial. Note that the number of trials in a sequence ranged between 4 and 12, and the number of search locations between 4 and 8 across experiments. Note further that the colors of each condition correspond to condition specific colors in subsequent plots. (**B-D**). Reaction times as a function of condition and trial position for (**B**) Experiment 1, (**C**) Experiment 2 and (**D**) Experiment 3. (**E**) Boxplot showing benefits of distractor location repetition in distractor-repeat sequences in Experiment 3 as a function of trial position contrasted to random distractor location repetitions in baseline sequences. Solid lines inside boxes show the mean, and dashed lines show the median.

### Experiment 1: The time-course of distractor learning

To study the neural mechanisms underlying learned distractor suppression, it is important to first characterize the rate at which suppression develops. We therefore first ran two behavioral experiments in which observers performed sequences of 12 visual searches with fixed distractor locations (Fig. 1A). In Experiment 1, in addition we included a baseline condition in which the target and the distractor location varied from trial to trial. The comparison between distractor-repeat and baseline sequences allowed us to assess the minimum number of distractor repetitions necessary to observe reliable suppression effects. In a second control condition, only a target was presented, the location of which varied over trials within a sequence to assess if, and if so when, learned suppression can completely counteract distractor interference. Critically, in this target-only variable condition, one of the placeholders was eliminated. In combination with a set size modulation (i.e., each sequence contained four or eight placeholders), this manipulation allowed us to exclude the possibility that the hypothesized suppressive effects in distractor-repeat sequences simply reflect facilitated processing at locations that are known to never contain a distractor. If so, distractor-repeat and target-variable conditions should demonstrate highly similar set size modulations.

#### Methods

Experiment and analyses were based on the *AsPredicted* registration at Open Science Framework (https://osf.io/t9qxu/).

##### Participants

A planned number of 18 participants (average (*M*) age = 24 years, range 19 – 34; seven men) participated in Experiment 1, in exchange for course credit or monetary compensation (10 euros per hour). Three participants were replaced, all because overall error rates were more than 2.5 SD’s below group average. Participants reported normal or corrected-to-normal vision and provided written informed consent prior to participation. The ethical committee of the Department of Psychology of the University of Amsterdam approved the study.

##### Apparatus, stimuli, design, and procedure

A Windows 7 PC running OpenSesame v3 (Mathôt, Schreij, & Theeuwes, 2012) using PsychoPy (Peirce, 2009) functionality generated the stimuli on an ASUS VG236 120-Hz monitor with a grey background, at ~80 cm viewing distance. Participants sat in a dimly lit room.

Each trial started with a 500-ms black fixation dot with a white rim (radius 5 pixels) followed by a placeholder display. This display contained four or eight black rimmed placeholders (radius 75 pixels), all placed on an imaginary circle centered (radius 225 pixels) on fixation. After a 250-ms delay, two Gabor patches (sf = 0.025, contrast = 1) were presented for 200-ms within the centers of two selected placeholders. One Gabor, the target, was tilted left or right (45° and 135°), while the other Gabor, the distractor, was randomly vertically or horizontally oriented. Participants had to indicate the orientation of the target Gabor (left or right) via keyboard response. They were instructed to respond as fast as possible, while trying to keep the number of errors to a minimum. A trial ended when participants made a response via a keyboard-press or 1000-ms after search display onset.

Crucially, there were three separate conditions. In the *distractor-repeat* condition, the location of the distractor was fixed across a sequence of 12 trials, while the location of the target was selected semi-randomly (i.e., the target location was never repeated on consecutive trials). In the *baseline* condition, target and distractor locations were random, with the restriction that neither the target and distractor location repeated on consecutive trials. In the *target-only variable* condition, the target was not accompanied by a distractor and its location was selected at random, again with the restriction that it did not repeat on consecutive trials. Also, in this condition, one of the placeholders was removed, in effect reducing the number of potential target locations by one relative to baseline and distractor-repeat conditions to establish whether repetition benefits in distractor-repeat sequences could be explained by shifts of attention away from the distractor location. At the start of each 12-trial sequence, the fixation point turned red for 50 ms. Search display set-sizes were fixed within repetition sequences.

Participants completed 36 practice trials in which only the distractor location was repeated, and 27 experimental blocks (9 blocks for each condition) of 72 trials each, with condition order counterbalanced over participants. Each experimental block contained three set size 4 and three set size 8 repetition sequences, randomly intermixed, resulting in 27 trials per trial position in the sequence for each condition. At the start of each block, participants were informed about the upcoming condition. Participants were encouraged to take a break between blocks.

##### Analysis

All data were analyzed in a Python environment (Python Software Foundation, https://www.python.org/). As preregistered, behavioral analyses were limited to reaction time (RT) data of correct trials only. RTs were filtered in a two-step trimming procedure: trials with reaction times (RTs) shorter than 200 ms were excluded, after which data were trimmed on the basis of a cutoff value of 2.5 SD from the mean per participant per condition. Remaining RTs were analyzed with repeated measures ANOVAs followed by planned comparisons with paired t-tests using JASP software (*JASP-TEAM, 2018*).

#### Results and discussion

##### Search times

Exclusion of incorrect responses (11.3%) and data trimming (1.8%) resulted in an overall loss of 13.1% of the data. Remaining RTs were entered into a repeated measures ANOVA with the within subject factors Condition (distractor-repeat, baseline, target-only variable), Set Size (4, 8) and Trial Position (1-12). As visualized in Figure 1B, the main effect of Condition (*F* (2, 34) = 99.1, *p* < 0.001) was accompanied by decreasing RTs over trials (*F* (11, 187) = 56,8, *p* < 0.001), and overall faster RTs at set size 4 than at set size 8 (*F* (1, 17) = 20.5, *p* < 0.001). An interaction between Condition and Trial Position showed that, as expected, decreasing RTs across trials was largely specific to distractor-repeat sequences (*F* (22, 374) = 12.142, *p* < 0.001), the only condition in which location expectations could develop. The overall slower response in the first compared to the second trial observed in all conditions, we attribute to the start of a new sequence.

Planned comparisons between baseline and distractor-repeat sequences demonstrated that a single repetition was sufficient to observe reliable benefits both at set size 4 (*t* (17) = 3.0, *p* = 0.008) and 8 (*t* (17) = 4.8, *p* < 0.001) and this benefit of distractor foreknowledge continued to increase across repetitions (all *t*’s > 3.2, all *p*’s < 0.006). However, distractor learning never completely counteracted distractor interference, as at the end of the sequence, RTs only numerically approached the target-only variable condition, but were still reliable slower (*t* (17) = 3.7, *p* = 0.002 for set size 4; *t* (17) = 5.6, *p* < 0.001 for set size 8).

A non-significant Set Size by Trial Position interaction suggested that the rate at which distractor learning developed was independent from the number of placeholders (*F* = 1.3, *p* = 0.22). Yet, as also visualized in Figure 1B and confirmed by a three-way interaction (*F* (22, 374) = 1.6, *p* = 0.041), distractor-repeat sequences did yield a significant Set Size by Trial Position interaction (*F* (11, 187) = 64.9, *p* < 0.001), which would also be expected if the repeated location was not suppressed, but observers instead shifted their attention to the remaining locations. To address this issue, we normalized the data relative to the first trial and fitted each individual’s data to an exponential decay function: D + (1 – D) * α^rep^, where D equals the absolute reduction relative to the first trial and *a* is rate at which this reduction develops. This analysis revealed that the set size modulation was evident in the reduction boundary (*t* (17) = 4.3, *p* < 0.001), but not in the learning rate (*t* = 0.2, *p* = 0.88). No such effects were observed in baseline or, importantly, target-only variable conditions (all *t*’s < 1.19, all *p*’s > 0.3), indicating that the observed benefits in distractor-repeat sequences reflect learned location suppression, rather than increased attention towards the remaining possible target locations.

To summarize, the results from Experiment 1 demonstrate that learning about the spatial probability of a distractor increases search efficiency. Distractor-based learning was implemented relatively quickly, although after 11 repetitions, distractor interference was not yet fully resolved, as RT was not yet as fast as in the target-only variable condition.

### Experiment 2: The time course of learned suppression and learned facilitation

Experiment 1 showed that 11 distractor location repetitions increasingly speeded up RTs, indicative of distractor learning. Although this effect differed between set sizes, it did not seem mediated by more attention to potential target locations (or a reduction in target location uncertainty), as the set size modulation was absent in target-only variable sequences and did not modulate learning rate in distractor-repeat sequences. Experiment 2 was designed to replicate the effect of distractor learning on performance observed in Experiment 1, and directly contrast this to the time-course of target learning.

#### Methods

The methods and analyses were preregistered (https://osf.io/t8c4m/) and were the same as in Experiment 1, except for the following changes: A planned number of 20 new participants (*M* = 23, range 18 – 27; seven men) participated. Two participants were replaced, one because overall reaction times were more than 2.5 standard deviations (SD’s) above group average, and one because overall error rates were more than 2.5 SD’s below group average.

In Experiment 2, there were two conditions. Next to the distractor-repeat condition (used in Experiment 1), here we also included a target-repeat condition, in which the location of the target was repeated over trials, while the location of the distractor was selected semi-randomly (i.e., the distractor location was never repeated on consecutive trials). All participants completed 18 experimental blocks, with repetition condition blocked in counterbalanced order.

#### Results and discussion

Exclusion of incorrect responses (10.8%) and data trimming (2.3%) resulted in an overall loss of 12.9% of the data. As visualized in Figure 1C, RTs again decreased across trials (*F* (11, 209) = 374.3, *p* < 0.001), with overall faster RTs in target than distractor repetition sequences (*F* (1, 19) = 106.4, *p* < 0.001) and for set size 4 compared to set size 8 displays (*F* (1, 19) = 34.0, *p* < 0.001). Significant interactions demonstrated that the repetition benefit was more pronounced when the target location was repeated (interaction between Condition and Trial Position; *F* (11, 209) = 24.3, *p* < 0.001) and when there were only four placeholders (interaction between Set Size and Trial Position; *F* (11, 209) = 2.2, *p* = 0.017). Critically, the effect of distractor repetition on RT was not differently affected by set size than the effect of target repetition on RT, as the three-way interaction between Condition, Trial Position and Set Size was not significant (*F* = 1.6, *p* = 0.11). Separate ANOVAs on the parameters resulting from the same model fits as in Experiment 1 yielded no significant Condition by Set Size interactions, neither on learning rate nor on reduction boundary (all *F*’s < 1.9, all *p*’s > 0.18), further arguing against the notion that in distractor-repeat blocks, observers simply paid more attention to potential target locations.

In summary, the results of Experiment 2 again demonstrate that distractor-based learning is implemented relatively quickly, although not as rapid as target-based learning. Together these experiments demonstrate that although a single repetition of the distractor already facilitates performance, the benefit of learning the spatial probability of a distractor increases gradually over trials. Next, we aimed to assess the neural mechanisms that underlie distractor learning-related behavioral benefits and contrast these to the neural mechanisms underlying target location learning: How does experience with distracting, but irrelevant information help the brain resolve distractor interference?

### Experiment 3: Neural mechanisms underlying learned facilitation and inhibition

Experiment 3 was designed to investigate the neural mechanisms that underlie the behavioral effects of distractor and target learning observed in the previous experiments. We were specifically interested to establish whether learning about the spatial probability of a distractor, just like learning about the spatial probability of a target, changes anticipatory tuning to the expected distractor location, reflective of preparatory inhibition, and/or suppresses distractor-related processing. To test this, we measured EEG and contrasted neural dynamics of learned suppression and facilitation across repetitions. We designed the experiment such that we could fit an IEM to reconstruct both the target and the distractor location from the multivariate EEG activity. This required that each location in the display was repeated equally often to prevent systematic biases in the model fits. Consequently, we had to reduce the sequence length to four repetitions, which based on our Experiments 1 and 2 was sufficient to observe robust behavioral benefits of distractor learning.

In addition, here, we aimed to parse out the influence of immediate distractor repetitions within the context of learned suppression. Based on the first two experiments, it remains unclear whether learned suppression goes above and beyond inter-trial priming. Previous studies have already shown that distractor repetition-related reductions in reaction time remain reliable when controlling for inter-trial priming effects in search tasks in which the distractor was more likely to occur at one location that remained the same across the entire experiment (Failing, Feldmann-Wüstefeld, Wang, Olivers, & Theeuwes, 2019; Wang & Theeuwes, 2018b). Yet, this control still allows for influences resulting from more distant trials in the past. Here, we investigated whether expected repetitions benefit distractor processing more than unexpected, random repetitions in a much shorter learning context.

#### Methods

The methods and analyses (preregistered at https://osf.io/4bx7y) were identical to those from Experiment 1 and 2, except for the following changes: A planned number of 24 new participants (*M* = 23, range 19 – 37; 6 men) participated in the experiment. Four participants were replaced: two because they did not complete all experimental sessions, and two because preprocessing (for details, see below) of EEG data resulted in exclusion of too many trials (> 30%). Participants were seated in a dimly lit testing room and all stimuli were presented at a distance of ~70 cm. Manual responses were collected via two purpose-built response buttons, which were positioned at the end of the armrests of the participant’s chair.

In contrast to Experiments 1 and 2, there were always six search locations and sequences were reduced to four trials. There were three experimental conditions, which were presented in separate blocks. In distractor-repeat and target-repeat conditions, the distractor or target location, respectively, was repeated. The location of the other item (target or distractor, respectively) was never repeated on consecutive trials and was placed maximally twice on the same location within a four-trial sequence. In the third, baseline condition, the target and distractor location were again variable with the restriction that the target location was never repeated and the distractor could maximally be presented two times at the same location in a sequence of four trials. This latter manipulation allowed us to contrast unexpected distractor repetition in baseline sequences (i.e., distractor priming) to learned suppression in distractor-repeat sequences. The length of the fixation display preceding each trial was randomly jittered between 450-ms and 750-ms, and each response was followed by a 200-ms blank screen.

Participants came to the lab twice. In each session, they completed 51 blocks of the task of 72 trials each, with condition blocked in counterbalanced order (e.g. target-repeat, baseline, distractor-repeat, etc), while their brain activity was recorded with EEG and their eye movements were monitored with an eye tracker. This resulted in 612 observations per condition and trial position within a four-trial sequence. In the first session, participants first completed a series of 24 practice trials in which only the distractor location was repeated.

##### Analysis software

Preprocessing and subsequent analyses were performed using custom written analysis scripts, which are largely based on functionalities implemented within MNE (Gramfort et al., 2014). These scripts can be downloaded at https://github.com/dvanmoorselaar/DvM.

##### EEG recording and preprocessing

EEG data were recorded at a sampling rate of 512 Hz using a 64-electrode cap with electrodes placed according to the 10-20 system (using a BioSemi ActiveTwo system; biosemi.com). All electrodes were rereferenced off-line to the average of two channels placed at the left and right earlobes respectively. External electrodes placed ~2 cm above and below the right eye and ~1 cm lateral to the external canthi were used to measure vertical (VEOG) and horizontal EOG (HEOG), respectively.

Continuous EEG of the two sessions was high-pass filtered using a zero-phase ‘firwin’ filter at 0.1 Hz as implemented in MNE to remove slow drifts, and subsequently epoched from −800 ms to 1300 ms relative to the onset of the placeholders, extended by 500 ms at the start and end of the epoch to control for filter artifacts during preprocessing and time-frequency analyses. The resulting epochs were baseline normalized using the whole epoch as a baseline. Prior to cleaning, EEG signals were visually inspected for malfunctioning electrodes, which were excluded from subsequent preprocessing steps (*M* = 2, range 0 – 5). To detect epochs contaminated with noise, we filtered the EEG signal with a 110 – 140 Hz bandpass filter and then used an adapted version of an automatic trial-rejection procedure as implemented in the Fieldtrip toolbox (Oostenveld, Fries, Maris, & Schoffelen, 2011) allowing for variable z-score cut-offs per participants based on the within-subject variance of z-scores (de Vries, van Driel, & Olivers, 2017). This resulted in an average rejection of 9.2% of all trials (range 2.3%–20.2%). After trial-rejection, ICA as implemented in MNE using ‘extended-infomax’ method was performed on non-epoched 1 Hz high pass-filtered data to identify and remove eye-blink components from the 0.1 Hz filtered data. Finally, malfunctioning electrodes were interpolated using spherical splines (Perrin, Pernier, Bertrand, & Echallier, 1989) before the data of the separate sessions was combined.

Throughout EEG recording, eye movements were monitored using an Eyelink 1000 (SR Research), sampled at 500 Hz. Gaze data were analyzed online, so that every time the participant broke fixation, auditory feedback was provided, signaling to the participant to keep their gaze at fixation. The Eyelink data was also analyzed offline. To control for drifts in the eye-tracker, epochs without a saccade (Nyström & Holmqvist, 2010) in the 300 ms pre-display interval were shifted towards fixation. Each epoch was then summarized by a single value indicating the largest deviation measured in a segment of data (> 40 ms). The results of the EEG analyses presented next are limited to trials with a fixation deviation <= 1° or in case of missing Eyelink data, trials with sudden sharp jumps in the HEOG as detected via a step algorithm (*M* = 7.2%, range = 0.3% - 18.3% combined for eye tracker analysis and step algorithm).

##### Inverted encoding model

To determine effects of distractor and target learning on (preparatory) spatial tuning of population-level activity, we applied an inverted encoding model (IEM; Brouwer & Heeger, 2009) to our EEG data to create spatial reconstructions of each location. This critically allowed us to examine effects of distractor and target learning on spatial tuning of population-level activity. The IEM procedure reverses the direction of inference from a decoding analysis by capturing the continuous relationship between stimulus position and multivariate EEG. Following Brouwer and Heeger (2009), we employed the same procedure reported in previous work (van Moorselaar et al., 2018) to separately reconstruct the target and the distractor location on the first and final trial across conditions.

Here, we modeled the hypothetical response in each of the six stimulus position channels (i.e., neuronal population) as half sinusoid raised to the seventh power and centered on the polar angle of each corresponding spatial channel. The resulting six basis sets were used to construct a *k* x *n* response prediction matrix C_1_, where *k* is the number of position channels (i.e. six) and *n* is the number of trials in the training set. To allow training and testing on independent data sets, the pre-processed EEG was partitioned into three sets, two training and one testing set. In doing so, we equated the number of observations across stimulus locations and conditions to prevent bias in the analysis. Consequently, a random subset of epochs was not included in data partitioning. To account for this, results were averaged across 10 data divisions, in which each of three data sets served as a testing set once. We then estimated the mapping from “channel” to “electrode” space by performing an ordinary least squares regression of the C1 matrix onto the *m* x *n_1_* observed power train matrix B_1_ where *m* is the number of electrodes (i.e. 64) and *n_1_* is the number of trials in the training set. This regression yields a *m* x *k* weight matrix W, where each electrode in *m* contains a regressor weight for each spatial channel in *k*. Next, in the test phase the model was inverted by performing ordinary least squares again, but now regressing these weights onto the *m* x *n_2_* observed power test matrix B_2_, where m is the number of electrodes and is the number of trials in the testing set. This regression transforms the test data B_2_ into an *k* x *n_2_* estimated channels responses matrix C_2_, where each spatial channel *k* contains an estimated response for all trials in n_2_. This procedure was repeated in a leave-one-out cross-validation routine until each block served as test set (i.e., B_2_). Finally, the estimated channel responses were averaged across trials in the testing set, separately for each trial type corresponding to the unique stimulus positions on the screen, and then shifted to a common center.

To evaluate reconstruction of the target and distractor location, we estimated channel tuning function (CTF) slopes. For this purpose, we collapsed across channels that were equidistant from the center and used linear regression to calculate the slope of the CTF, where a positive slope indicates greater location selectivity (relative to the other locations), a flat slope no location specificity, and a negative slope reduced location selectivity (relative to the other locations). Previous work using IEMs has demonstrated that the reconstruction of spatially selective CTFs during preparatory spatial attention is largely specific to power in the alpha band (Foster et al., 2017; Samaha et al., 2016; van Moorselaar et al., 2018). To isolate frequency-specific activity, we band-pass filtered the data using a fifth-order butterworth bandpass filter within MNE. After filtering, the EEG signal was down-sampled to 128 Hz to reduce computational time of the IEM analysis. To confirm that reconstruction was most pronounced within the alpha band, we first searched a broad range of frequencies (4-30 Hz, in increments of 2Hz with a 4-Hz band; 4-8 Hz, 6-10 Hz, etc) using the collapsed data of all conditions of interest. Total power serving as input for matrix B_1_ was then calculated after extraction of the complex analytic signal via a Hilbert transform and then averaged across trials, such that power reflects the ongoing activity irrespective of its phase relationship to the onset of the stimulus displays. The IEM routine was subsequently applied to each time-frequency point. The same analysis was also performed using evoked power, which was calculated by averaging the complex analytical signal across trials before power extraction, such that evoked power reflects activity phase-locked to stimulus onset.

##### Event-related potential (ERP) analysis

We also examined how learning changed distractor and target processing after stimulus presentation using ERPs. To enable isolation of target- and distractor-specific ERP components, the primary analysis focused on ERPs elicited by lateralized targets presented below the horizontal midline when the distractor was presented on the midline, and vice versa. Waveforms were computed separately for ipsilateral and contralateral scalp regions.

After cleaning, epochs were 30 Hz low-pass filtered and baseline corrected using a −300 to 0 ms pre-placeholder onset baseline period. ERP averages were balanced to contain an equal number of observations across conditions and trial position. We were primarily interested in ERP components related to visual processing and attention, namely the P1, N1, N2pc and the Pd, which are typically observed at lateral posterior electrodes sites. Based on visual inspection of the topographic distribution of condition-averaged voltage values in these regions, we selected O1/O2, PO3/PO4, and PO7/PO8 as our electrodes of interest.

P1 and N1 windows, respectively 110 - 150 ms and 160 - 220, were selected based on visual inspection of the group and condition-averaged waveforms. For each subject, we obtained the mean voltage value over a 35-ms time window centered around the P1 or N1 peaks within these selected time windows. N2pc and Pd windows, respectively 170 - 230 ms and 280 – 360, were also selected via visual inspection of the group and condition-averaged waveforms, but now using contralateral – ipsilateral difference waveforms.

##### Backward Decoding Model (BDM)

Finally, we also applied multivariate decoding analysis (MVPA) to decode target and distractor locations. This approach allowed us to examine whether the neural signal contained location-specific information that may not have been captured by the assumptions in our inverted encoding model. Also, this approach provides a more fine-grained spatial profile of attentional selection than lateralized ERP components (Fahrenfort, Grubert, Olivers, & Eimer, 2017), which by definition are driven by activation differences between cortical hemispheres.

To decrease the computational time of the decoding analysis, the EEG signal was down-sampled to 128 Hz. We applied a linear classification algorithm (Pedregosa et al., 2011) in electrode space (64 electrodes) at each time sample, using a 10-fold cross validation scheme. Per condition and trial position (first, last) the dataset was randomly split into 10 equally sized subsets, while ensuring that each decoding label (i.e., spatial location; 1-6) was selected equally often. Subsequently, the classifier was trained on 90% of the data and tested on the remaining 10% of the data. Training and testing on independent datasets was repeated until all data was tested exactly once.

##### Statistics

To evaluate how location repetition changed the neural representation of target and distractor locations we contrasted target-repeat sequences and distractor-repeat sequences with baseline sequences using group-level permutation testing with cluster correction for multiple comparisons (Maris & Oostenveld, 2007). First, for the first and final trial within a sequence, the conditions of interests were contrasted to the baseline condition (e.g., target-repeat 4 vs. baseline 4). Second, significant clusters on the final trial were evaluated again after controlling for potential differences between repeat and baseline sequences at the start of the sequence (e.g., target repeat 4 - baseline 4 vs. target repeat 1 - baseline 1).

In addition to cluster-based group level permutation testing, specific comparisons were also evaluated using repeated measures ANOVAs. While permutation testing has the merit of not choosing specific time points a priori and therefore, allows to observe potentially non-expected effects, it also has the risk of missing brief, but reliable effects. Moreover, as tests are conducted per sample, they are more sensitive to noise than analyses conducted on measures that are obtained by averaging across several samples. Therefore, when we expected ERP effects within specific time-windows, we also evaluated their presence using repeated measures ANOVA’s, as is common in ERP research. These tests were also used in case only one of the two permutation contrasts described above identified a significant cluster.

## Results

### Search times

Exclusion of incorrect responses (7.2%) and data trimming (2.4%) resulted in an overall loss of 9.7% of the data. Figure 1D illustrates that RTs decreased with repetition (main effect Trial Position; *F* (3, 69) = 105.1, *p* < 0.001), with overall fastest RTs following target repetition and slowest RTs without repetition (main effect Condition; *F* (2, 46) = 60.8, *p* < 0.001). While repetition benefits were most pronounced in target-repeat blocks (interaction effect Condition and Trial Position; *F* (6, 138) = 101.2, *p* < 0.001), distractor repetition also speeded search: A separate ANOVA without target-repeat blocks also yielded a highly significant interaction effect (*F* (3,69) = 54.5, *p* < 0.001). Planned pairwise comparisons demonstrated that within both target and distractor repeat sequences each subsequent repetition speeded RTs (all *t*’s > 4.0, all *p*’s < 0.001), whereas no such effect was observed in the baseline condition in which targets and distractor occurred at different locations from trial-to-trial and no expectations could develop (all *t*’s < 0.4, all *p*’s > 0.7). These observations show that both benefits of target and distractor learning developed gradually across repetitions. Importantly, as visualized in Figure 1E, the first repetition in distractor-repeat sequences speeded search more than random repetitions in baseline sequences (*t* (23) = 2.5, *p* = 0.021). This finding indicates that although a part of our behavioral effects was likely driven by inter-trial priming, learned biases are additive to these pure repetition effects, in line with previous reports (Failing et al., 2019; Wang & Theeuwes, 2018b)

### Inverted encoding model (IEM) analysis

To establish whether the observed benefits of location repetition may be brought about by preparatory biasing of visual regions representing the expected location, we used an IEM to reconstruct the target and distractor location from the pattern of EEG data. Previous research has demonstrated that these models are sensitive to the attended location during the interval preceding stimulus onset, in line with the notion that attention can prioritize target processing by sharping spatial tuning to the task-relevant location in advance (Foster et al., 2017; Samaha et al., 2016). We thus expected target learning in our study to be associated with enhanced reconstruction of the target location before search display onset. Yet, the main question was if and how, distractor learning also changes the neural representation of the distractor location in anticipation of distracting input.

Based on previous research (Foster et al., 2017; Samaha et al., 2016; van Moorselaar et al., 2018), we expected any effects to be especially present in the alpha-band. An analysis collapsed across conditions and repetitions across a range of frequencies confirmed that sustained reconstruction of both the target and the distractor location was especially pronounced within the alpha-band (8 - 12 Hz; Fig. 2). Having established that topographic distribution of alpha power tracked both the location of the target and the distractor, we examined how CTF reconstruction was modulated by location repetition within the alpha band.

**Figure 2.**
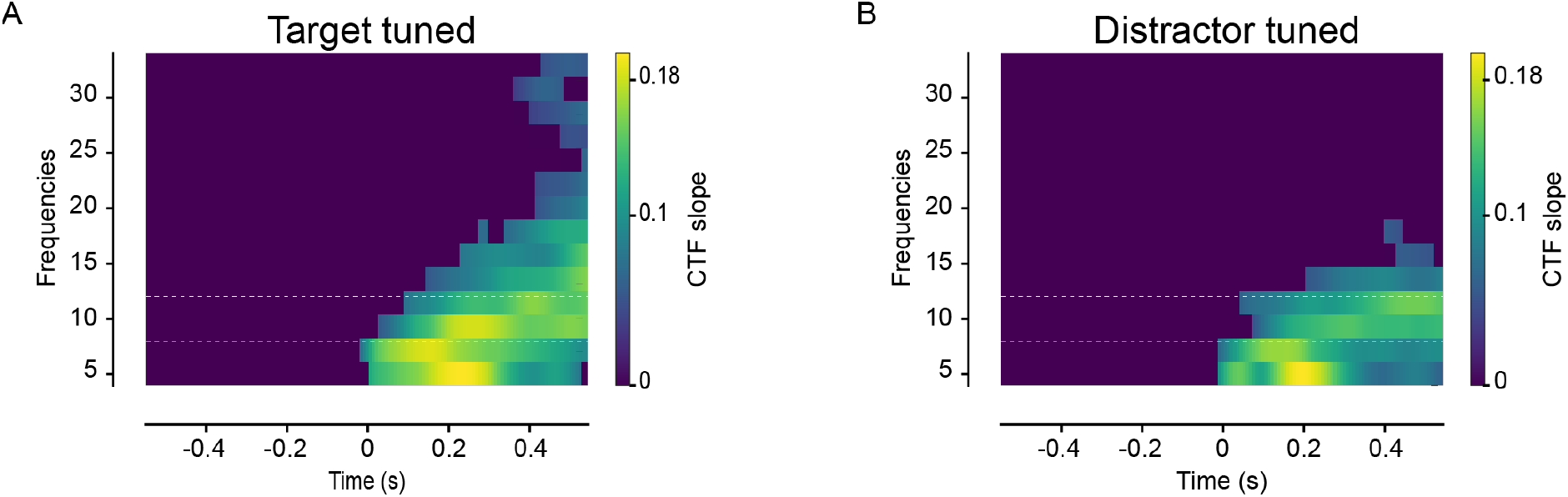
Topographic power in a range of low frequencies tracks both the location of the target and the distractor. (**A**) Total power CTF slopes tuned to the target location across a range of frequencies and collapsed across all conditions of interest (i.e. DvTv1, DvTv4, DvTr1, DvTr4). All non-significant values were set to zero in a two-step procedure. First, each individual data point was tested against zero with a paired-sampled t-test. After setting non-significant values to zero data were evaluated using cluster-based permutation. (**B**) Total power CTF slopes tuned to the distractor location across a range of frequencies and collapsed across all conditions of interest (i.e. DvTv1, DvTv4, DrTv1, DrTv4).

### Target CTF

Figure 3 shows the reconstruction of the target location in baseline and target-repeat blocks for the first and final trial in a four-trial sequence using either evoked (phase-locked) (Fig. 3A) or total (Fig. 3B) alpha power. Target location repetitions increased spatial tuning immediately following search display onset both within evoked and total power. Within evoked power, the identified cluster remained reliable after controlling for any differences on the first trial of the sequence (grey bars in Fig. 3A; right plot). Consistent with previous findings (Foster et al., 2017; Samaha et al., 2016), only total alpha-power enabled reconstruction of the target location in anticipation of display onset, confirming that target location learning sharpened spatial tuning to the target location at the neural level in advance (Fig. 3B; right plot). This anticipatory effect became weaker and statistically insignificant after the onset of the (empty) placeholders, suggesting that their onset may have temporarily obscured the anticipatory tuning. The observed significant target repetition-related anticipatory tuning did not survive the cluster-based baseline correction. We therefore explored this anticipatory tuning further using a repeated measures ANOVA on the average tuning slopes within the pre-stimulus period (−550 ms – 0 ms), which critically yielded a significant interaction between Condition (baseline, target repeat) and Trial Position (1,4) (*F* (1,23) = 7.0, *p* = 0.013).

**Figure 3.**
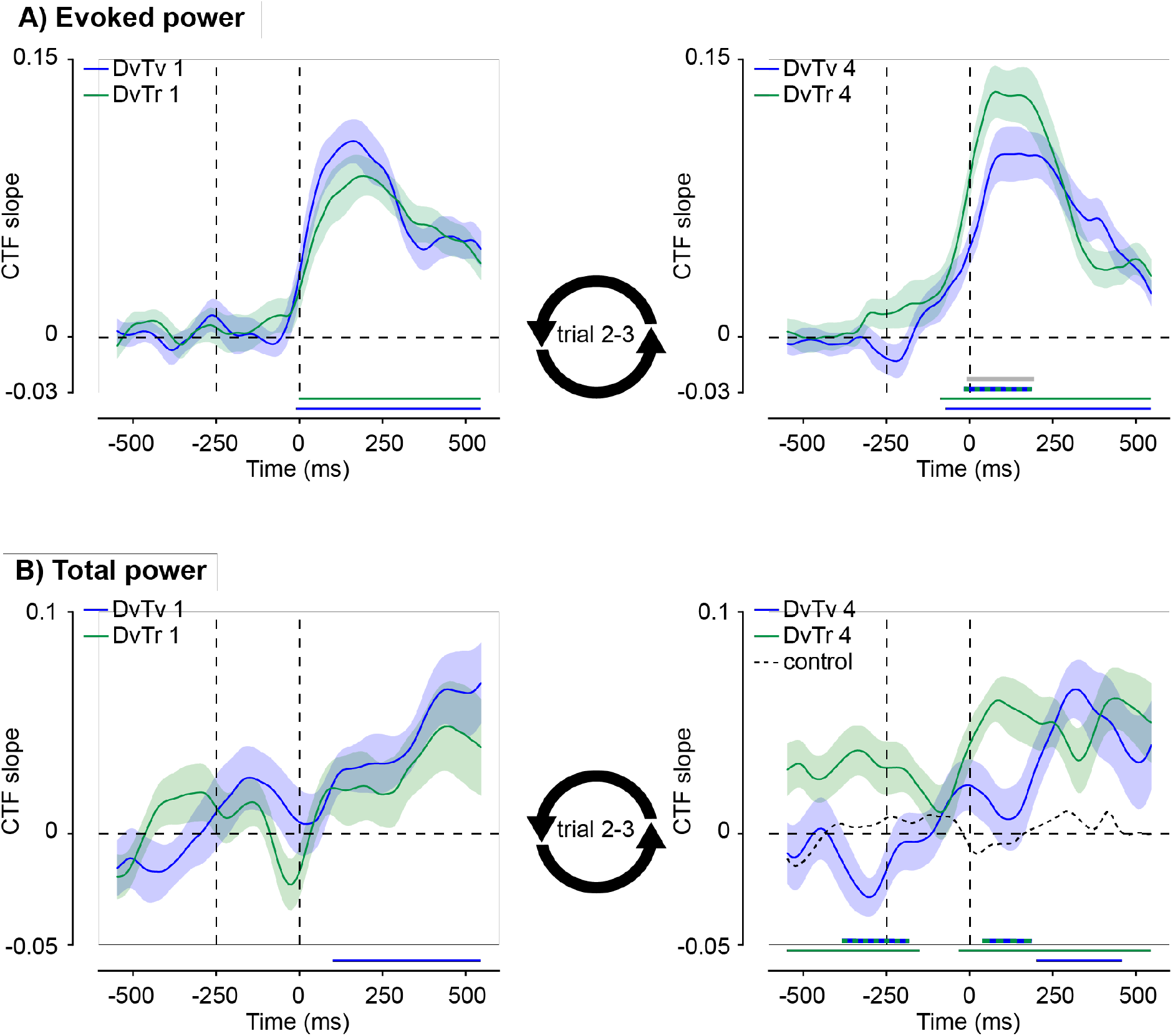
Target repetition increased anticipatory and post-stimulus spatial tuning to target locations. All plots show the CTF slope, which here quantifies the location specificity of the topographic distribution of activity in the alpha band. (A) Evoked power CTF slopes tuned to the target location at the first (left plot) and final (right plot) trial in baseline (DvTv) and target-repeat sequences (DvTr). (B) Total power CTF slopes tuned to the target location at the first (left plot) and final (right plot) trial in baseline and target-repeat sequences. Target repetition increased spatial tuning to the predictable target location already in advance of target presentation. A control (dotted black line) analysis showed that this effect cannot be attributed to lingering effects from the preceding trial. Shaded error bars reflect bootstrapped *SEM* (same applies to subsequent figures). Colored bars on the x-axis (blue; green) indicate time points where conditions differ significantly from 0 after cluster correction (p <0.05). Double-colored thick lines indicate time points with a significant difference between the respective conditions after cluster correction (p <0.05).

To exclude the possibility that lingering target processing from the preceding trial may have contributed to the observed increase in anticipatory tuning in target-repeat blocks, in a control analysis, we replaced the location labels in the baseline condition with those from the preceding trial in the sequence. If the observed anticipatory tuning following target repetition was driven by lingering activations elicited by target selection on the preceding trial, this analysis should also reveal anticipatory tuning. However, as shown in Fig.3B (right plot; dotted black line), no such tuning was observed. Together, these results indicate that target foreknowledge was associated with enhanced tuning of spatially selective neural populations to the target location already in anticipation of the search display, and this spatial selectivity increased further in response to visual input.

### Distractor CTF

We next examined how distractor learning affected the neural representation of the distractor location. Reconstruction of the distractor location, as shown in Figure 4, showed a markedly different pattern. Both within evoked (Fig. 4A) and total alpha power (Fig. 4B) we observed no reactive modulation of spatial tuning by distractor location repetition, nor were there any reliable changes in anticipatory tuning to the distractor location. Reconstructions using total power appeared to become more negative as a function of distractor repetition around search display onset. Yet, this negative tuning was non-significant nor did it differ from the baseline condition, suggesting that this may have been an artifact of our analysis and/or a highly variable effect across subjects.

**Figure 4.**
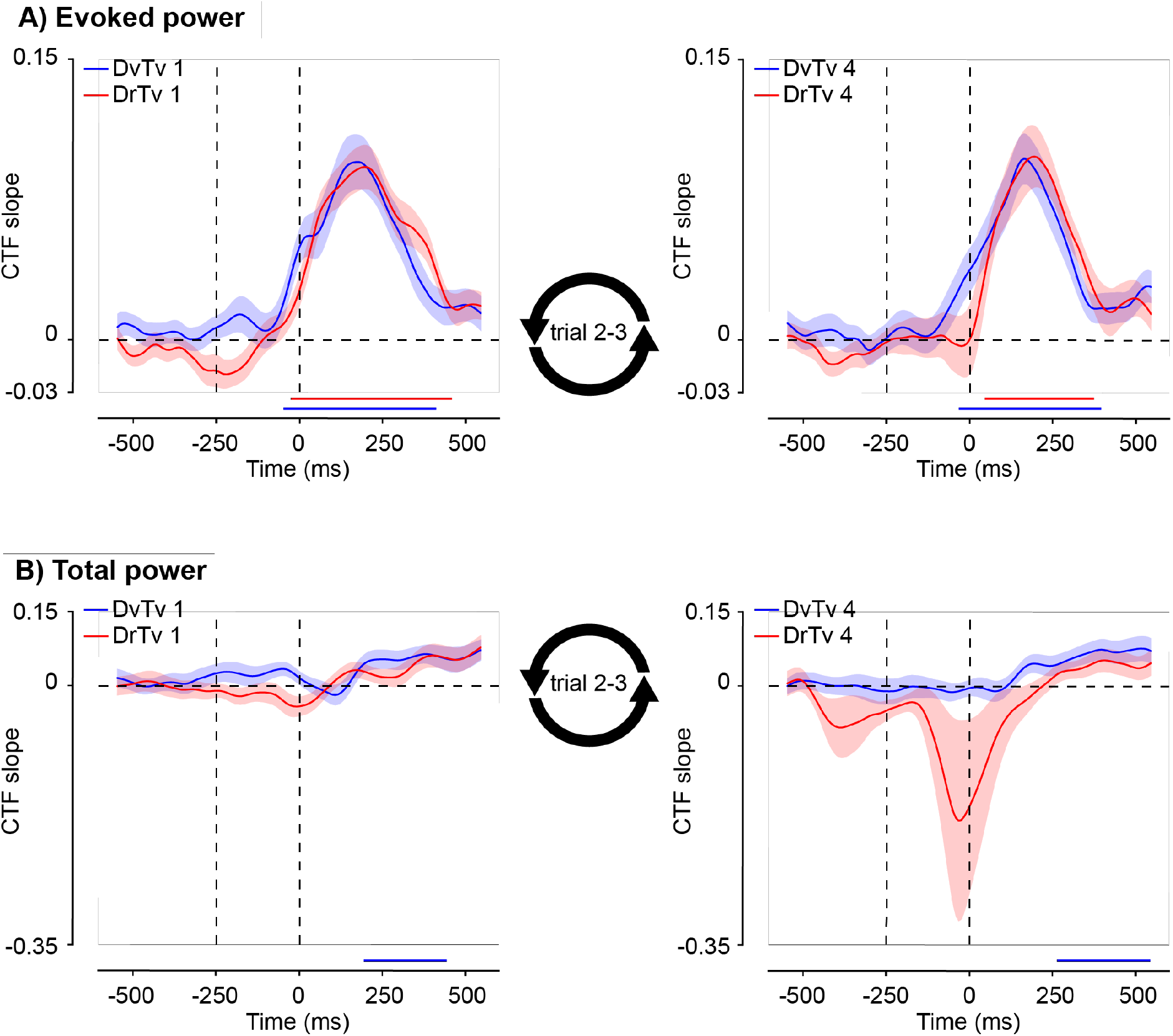
Distractor repetition did not change spatial tuning to distractor locations. (**A**) Evoked alpha power CTF slopes tuned to the distractor location at the first (left plot) and final (right plot) trial in baseline (DvTv) and distractor-repeat sequences (DrTv). (**B**) Total alpha power CTF slopes tuned to the distractor location at the first (left plot) and final (right plot) trial in baseline and distractor-repeat sequences. Colored bars on the x-axis (blue; red) indicate time points where conditions differ significantly from 0 after cluster correction (p <0.05).

Thus, while target repetition was associated with both increased anticipatory and post-stimulus spatial tuning, spatial tuning to the distractor location appeared unaffected by distractor repetition. This dissociation was confirmed by a cluster-based permutation comparison between distractor-repeat and target-repeat, which yielded both an anticipatory (−338 ms till −167 ms) as well as a reactive significant cluster (−6 ms till 264 ms; p < 0.05). This finding suggests that the observed reduction in distractor interference at the behavioral level is not mediated by changes in the neural representation of the distractor location, as measured with EEG and CTFs.

### Event-related potential (ERP) analysis

To determine whether repetition changed early visual processing, we examined the amplitude of the visual-evoked potentials, P1 and N1. Modulations of the amplitude of these exogenous components by attention-directing spatial cues have been linked to different aspects of attention, with the P1 effect reflecting attentional inhibition and the N1 effect signaling attentional enhancement (Couperus & Mangun, 2010; Freunberger et al., 2008; Luck et al., 1994; Slagter, Prinssen, Reteig, & Mazaheri, 2016). In addition, we evaluated how location learning affected later-stage target and distractor processing, as reflected in the lateralized ERP components, the N2pc and the Pd, respectively. Whereas the N2pc is thought to reflect attentional selection (Eimer, 2014; Luck, 2012; Luck & Hillyard, 1994), the Pd is selectively elicited by distracting visual input and linked to distractor inhibition (Gaspelin & Luck, 2018a; Hickey, Di Lollo, & McDonald, 2009).

### Target ERPs

Contrary to our expectations, cluster-based permutation tests identified no significant effects of target repetition within the P1 or N1 time window. To further explore the effects of target repetition, mean P1 and N1 voltages were entered into separate repeated measures ANOVA’s with the within subjects factors Condition (variable vs. repeat), Trial Position (1 vs. 4) and Laterality (contra vs. ipsi). This analysis also revealed no Condition by Trial Position interaction (*F* = 2.6, *p* = 0.12 for P1; *F* = 3.1, *p* = 0.09 for N1), suggesting that early visually evoked potentials were unaffected by target repetition.

By contrast, as visualized in Fig. 5, target repetition did reduce the N2pc, an ERP index of stimulus-driven attentional orienting, as captured by a significant baseline-corrected cluster within the typical N2pc time window. As the direct comparison on the final repetition between baseline and the target repeat condition using cluster-based permutation testing revealed no significant cluster, we also explored the N2pc modulation with a repeated measures ANOVA with N2pc amplitude values averaged over the N2pc time window as the dependent variable. This yielded a significant three-way interaction (*F* (1,23) = 10.4, *p* = 0.004), reflecting a reduction of the N2pc amplitude with target location repetition (*t* (23) = 3.7, *p* = 0.001), whereas this reduction was absent in baseline trials without repetition (*t* = 0.5, *p* = 0.62). This set of findings again illustrates enhanced statistical power of first averaging over samples within a specific time window and conducting statistical analyses on this average versus a cluster-based permutation test on individual samples, which is conceivable more sensitive to noise. An exploratory jack-knife procedure (Miller, Patterson, & Ulrich, 1998) demonstrated that the N2pc returned earlier to baseline at the final relative to the first target repetition (Δ = 28 ms, *t* = 3.38).

**Figure 5.**
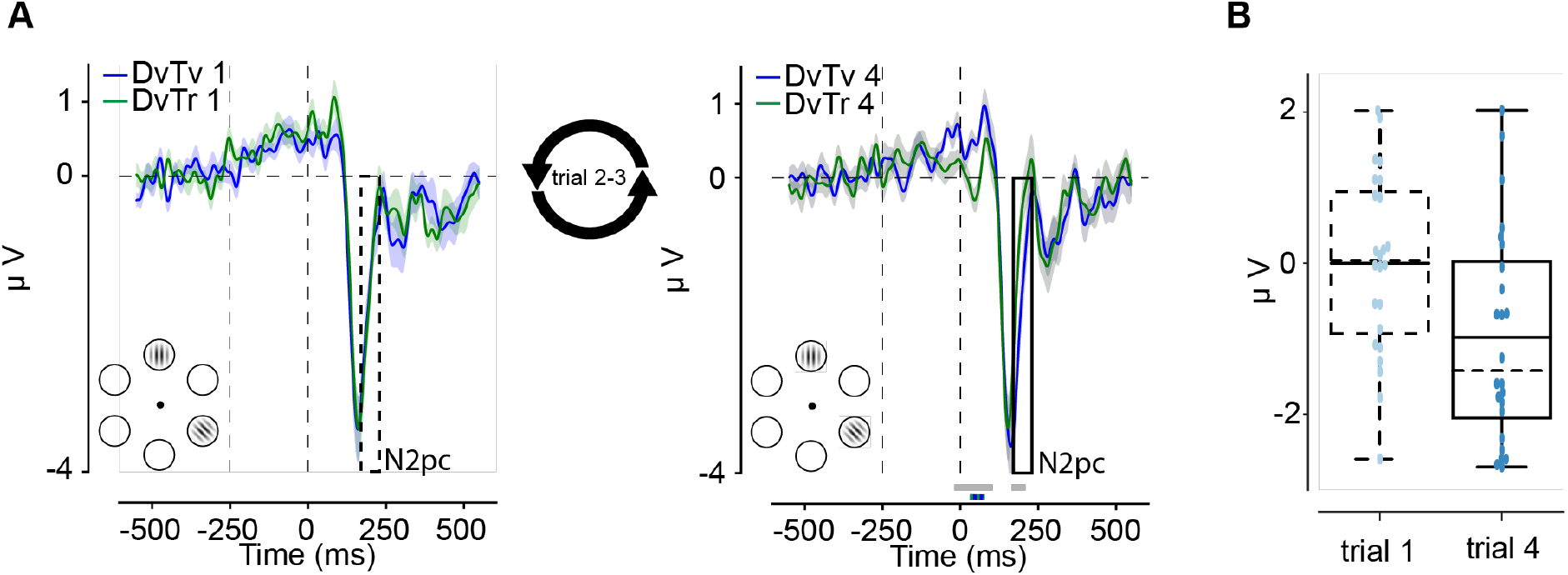
Target repetition reduced the amplitude of the target-evoked N2pc. ERPs evoked by targets were computed only using trials where the target was presented on the bottom left or right from fixation, with a distractor on the midline. (**A**) Difference waveforms (contralateral – ipsilateral) revealing the N2pc are shown separately for the baseline (DvTv) and target-repeat (DvTr) condition on the first (1) and final (4) repetition in the sequence. Double-colored thick lines indicate time points with a significant difference between the respective conditions after cluster correction (p <0.05). Grey thick lines indicate time points with a significant condition difference after baseline correction (p <0.05). (**B**) Boxplots show the difference between conditions (DvTv – DvTr) on the first (dashed) and final (solid) trial of a target repetition block within the N2pc window (170-230 ms). Solid lines inside boxes show the mean, and dashed lines show the median.

### Distractor ERPs

ERP waveforms evoked by the distractor yielded a different pattern of results. Again, contrary to our predictions, we observed no significant effects within the P1 or N1 time windows (*F* = 0.0, *p* = 0.99 for P1; *F* = 0.8, *p* = 0.37 for N1). Like targets, distractors elicited a clear N2pc (*F* (1,23) = 46.9, *p* < 0.001) indicating that distractors captured attention on at least a subset of trials, but notably, in contrast to target repetition, this component was not modulated by distractor repetition. No significant clusters were observed within the N2pc window, nor yielded the repeated measures ANOVA any significant interactions (all *F*’s < 1.11, all *p*’s > 0.3). At the first trial of a sequence, distractor N2pc’s, however, were followed by a positivity, a positivity that was absent in target-elicited waveforms, and furthermore, given its latency and scalp topography is likely to be the Pd, an ERP related to distractor inhibition (Gaspelin & Luck, 2018b). Interestingly, as shown in Fig. 6, this positivity was greatly reduced in the final distractor repetition trial. A significant cluster obtained when contrasting baseline and distractor-repeat at the final repetition confirmed that the amplitude of this positivity reduced with distractor repetition. As the observed reduction failed to survive the cluster-based permutation baseline correction, Pd amplitudes were also evaluated via a repeated measures-ANOVA. A significant threeway interaction (*F* (1,23) = 4.8, *p* = 0.038) and post-hoc paired t-tests, confirmed that the reduction of the Pd amplitude was specific to distractor repetition condition (*t* (23) = 2.9, *p* = 0.007), whereas it was absent in the baseline condition (*t* = 0.78, p = 0.44).

**Figure 6.**
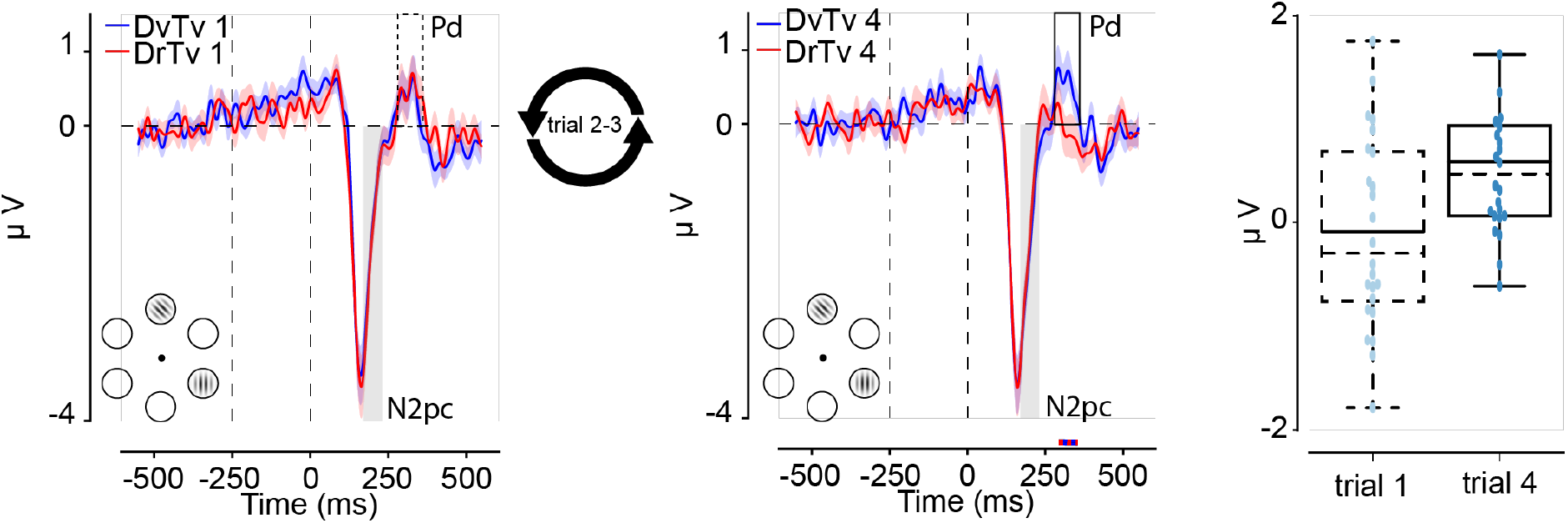
Distractor repetition reduced the distractor-evoked Pd. ERPs evoked by distractors were computed based on trials where the distractor was presented on the bottom left or right from fixation, with a target on the midline. (A) Difference waveforms (contralateral – ipsilateral) revealing the N2pc and Pd are shown separately for the baseline (DvTv) and distractor-repeat (DrTv) conditions on the first (1) and final (4) repetition in the sequence. Double-colored thick lines indicate time points with a significant difference between the respective conditions after cluster correction (p <0.05). (B) Boxplots show the difference between conditions (DvTv – DrTv) on the first (dashed) and final (solid) repetition within the Pd window (280-360 ms). Solid lines inside boxes show the mean, and dashed lines show the median.

Thus, ERP analyses showed that target repetition resulted in an earlier offset the N2pc, an ERP component associated with attentional selection, while distractor repetition selectively modulated the amplitude of the Pd, an ERP component associated with inhibition of distracting visual input. Against our predictions, however, we observed no evidence that early visual processing, as reflected in the P1 or the N1, was modulated by target or distractor location foreknowledge. Possibly, overlapping P1/N1 ERPs evoked by the other stimuli in the display masked effects by location foreknowledge on these early visual-evoked ERPs.

### Multivariate decoding: target location

The classification accuracy results in Figure 7 show that the decoding model could discriminate the target location, but in contrast to IEM, only after search display onset. Moreover, target repetition reliably modulated decoding classification accuracy. Specifically, mimicking the N2pc findings, the classification peak narrowed within the N2pc time window, as reflected by a significant baseline-corrected cluster (Fig 7). That is, decoding accuracy more quickly reduced after target learning, suggesting that target repetition may have reduced the duration of time the target location was represented in this time window. In contrast to our IEM results, decoding did not provide evidence for target learning-induced changes in the spatial representation of the target location prior to target presentation. To exclude the possibility that anticipatory decoding was obscured by the broad band of frequencies used, we repeated the same analysis with alpha-band filtered data (as in our IEM analysis) using either all or a set of 32 posterior electrodes. However, this exploratory analysis also only revealed post-stimulus modulation of target decoding by repetition, but no anticipatory effects. Thus, the decoding results more closely followed the univariate ERP analysis results.

**Figure 7.**
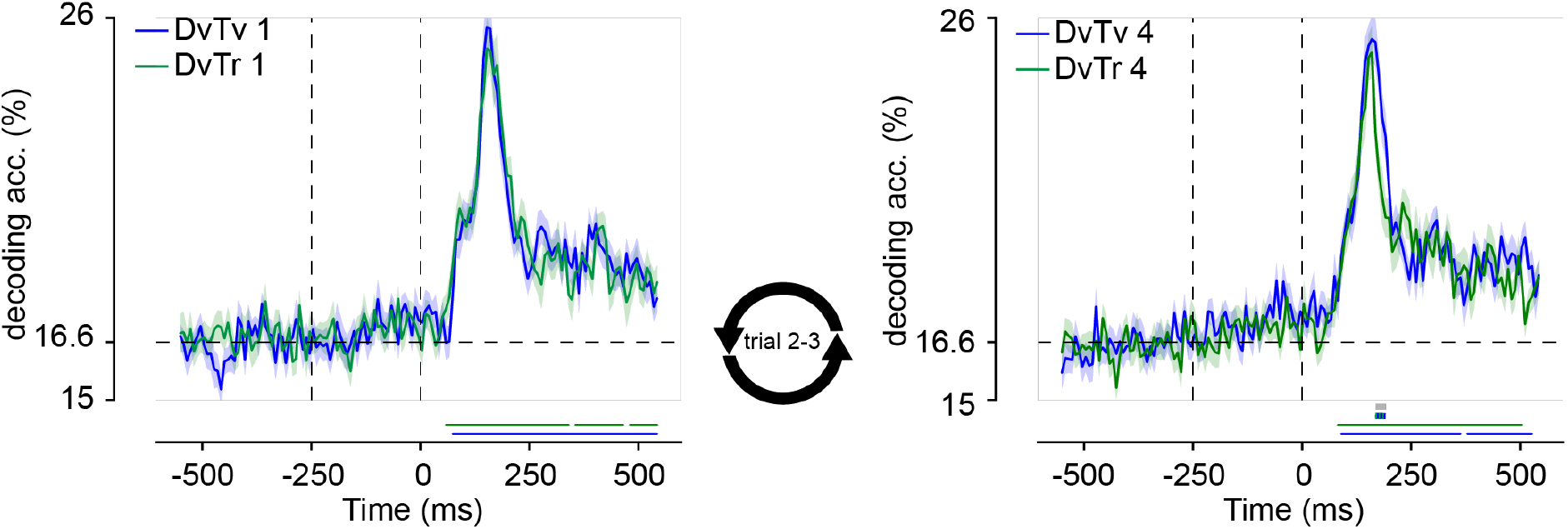
Target repetition was associated with a shortening of the representation of the target location within the N2pc time window, as reflected in decoding accuracy. Shown are target-location decoding accuracies of broad-band EEG using all 64 electrodes separately for baseline (DvTv) and target-repeat (DvTr) sequences and the first (1) and last (4) trial in a sequence. Colored bars on the x-axis (blue; green) indicate time points where conditions differ significantly from 0 after cluster correction (p <0.05). Double-colored thick lines indicate time points with a significant difference between the respective conditions after cluster correction (p <0.05). Grey thick lines indicate time points with a significant condition difference after baseline correction (p <0.05).

### Distractor decoding

We again observed a different pattern for distractor location decoding than target location decoding. Broadband EEG decoding was also sensitive to the distractor location (Fig. 8), and only post distractor, but distractor expectations did not appear to modulate classification accuracy. If anything, the classification peak within the N2pc window appeared to narrow, but this was not confirmed by cluster-based permutation tests. However, an exploratory repeated measures ANOVA yielded a significant interaction between Condition and Trial Position (*F* (1, 23) = 12.6, *p* = 0.002), indicating that distractor location representation may have actually been modulated by distractor location repetition in the time window of the N2pc, as was also observed for the target location as a function of target location repetition.

**Figure 8.**
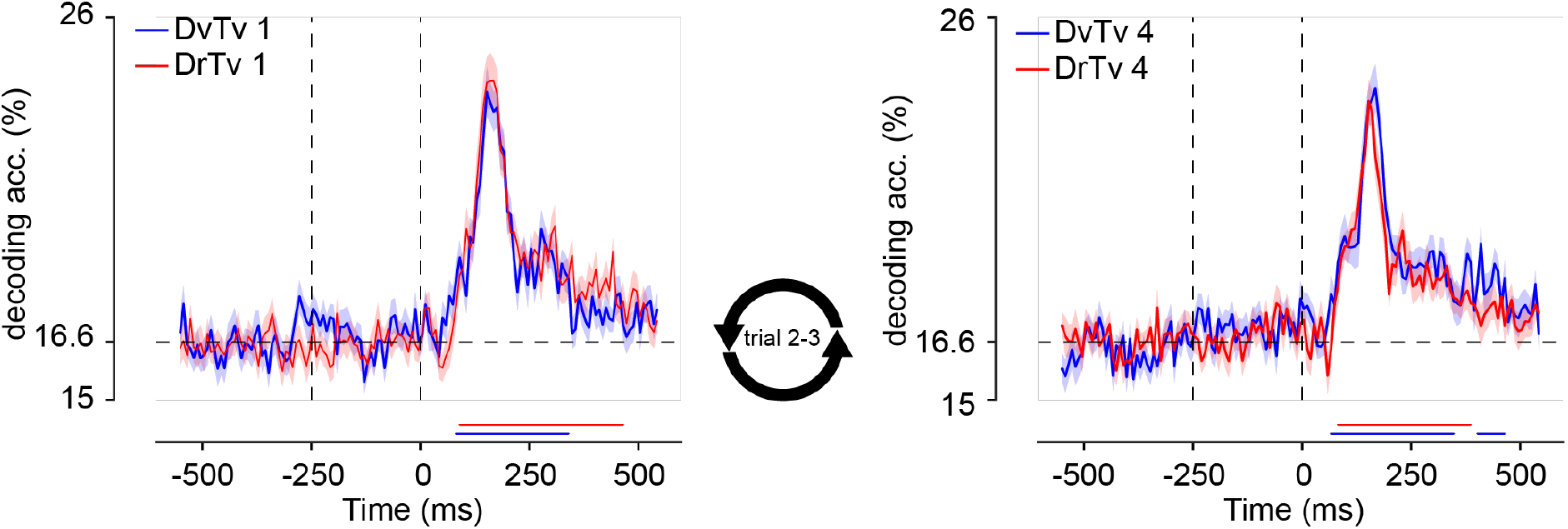
Distractor repetitions was not associated with a change in the distractor representation, as reflected in decoding accuracy. Shown are distractor-location decoding accuracies of broad-band EEG using all 64 electrodes separately for baseline (DvTv) and distractor-repeat (DrTv) sequences and the first and last trial in a sequence. Colored bars on the x-axis (blue; red) indicate time points where conditions differ significantly from 0 after cluster correction (p <0.05). Double-colored thick lines indicate time points with a significant difference between the respective conditions after cluster correction (p <0.05).

## General Discussion

The aim of the present study was to gain more insight into how the brain learns to ignore distracting information based on past experience, and to what extent these mechanisms differ from learning about relevant aspects of the environment. The benefits of distractor learning, which developed at a slower rate than target learning, could not be explained by facilitation of non-distractor locations and it could not be explained by priming alone. In line with recent findings (Foster et al., 2017; Samaha et al., 2016) and in support of top-down facilitation, we found that expectations about the upcoming target location sharpened neural tuning to that location before target presentation, and furthermore, reduced post-target processes related to attentional target selection (Eimer, 2014; Luck, 2012), as indicated by the N2pc ERP component. In contrast, expectations about the upcoming distractor location only modulated post distractor processing, i.e., reactively. Most notably, expected distractors appeared to no longer elicit a Pd, an ERP component related to distractor inhibition (Gaspelin & Luck, 2018b; Hickey et al., 2009), as if distractors were no longer considered distractors by the brain. Together, these findings suggest that target facilitation and distractor suppression are differently influenced by learning, and thus, at least in part, rely on different neural mechanisms. More generally, our results demonstrate the complementary nature of findings obtained with forward encoding modeling, univariate ERP analysis, and multivariate decoding, as these different types of analyses were in part sensitive to different aspects of the data, as discussed in more detail below.

Spatial tuning to the expected target location within the alpha-band already emerged in the interval preceding visual search, likely reflecting greater top-down attention to that location (Foster et al., 2017; Samaha et al., 2016). Strikingly, no changes in spatial tuning were observed to expected distractor locations, despite clear behavioral benefits of distractor location foreknowledge and a substantial reduction in the need for reactive distractor inhibition, as captured by a much smaller distractor-evoked Pd. These findings corroborates findings from a recent study by Noonan et al. (2016) that did not observe evidence for a role of alpha oscillations in preparatory distractor suppression, and also used a visual search task with more than 2 locations, preventing participants from simply directing more attention to the target location. Together, these findings call the computational role of alpha oscillations into question (Foster & Awh, 2018). One dominant view is that activity in sensory regions representing the distracting information can be top-down inhibited in advance through increasing alpha-band activity (Foxe & Snyder, 2011; Jensen & Mazaheri, 2010), just like knowledge about upcoming targets can be used to increase the excitability of task-relevant sensory regions to prioritize target processing by releasing inhibition by alpha oscillations. Our findings and the Noonan et al. (2016) findings do not provide support for the notion that distractor inhibition is implemented through preparatory inhibition of activity in visual regions representing the distractor location. While one should always be careful when interpreting a null finding, our results are more in line with a predictive coding framework in which processing of any expected stimulus is suppressed because it provides little new information (Noonan et al., 2018). Specifically, we found that learning about the upcoming distractor location greatly reduced the Pd component, which was selectively elicited by distractors before learning and is considered a neural marker of distractor inhibition (Gaspelin & Luck, 2018b; Hickey et al., 2009). Previous research has associated faster RTs with larger Pd amplitudes (Gaspar & McDonald, 2014) and shown that salient distractors that fail to capture attention elicit a Pd, indicative of active inhibition (Gaspelin & Luck, 2018a). Yet, here we found that distractor learning-related improvements in performance were associated with a *reduction* in Pd amplitude. This finding may suggest that when a distractor is expected, there is no more need for active inhibition, because the brain has learned that it can be safely ignored.

Further supporting the idea that target facilitation and distractor suppression rely on different neural mechanisms, we observed that, in contrast to distractor repetition, the N2pc amplitude by repeated targets was selectively reduced (see Praamstra, 2006 for a similar finding). Specifically, a jack knife procedure revealed the N2pc returned to baseline 28ms earlier in the last vs. the first trial within a target repetition sequence. This could indicate that post-stimulus attentional selection was more quickly resolved. Alternatively, in line with the idea that the N2pc signals an object individuation process (Mazza & Caramazza, 2015), target individuation may have been more efficiently resolved when the target appeared at the predicted location and therefore was more precisely represented. By contrast, the N2pc elicited by distractors appeared to be insensitive to repetition effects, indicating that distractors continued to capture attention to the same extent. Distractors may have captured attention because the target location was random and attention thus had to be guided via a feature template representation in memory (left or right tilted gabor), which by design was highly similar to the distractor (horizontally or vertically oriented gabor). Previous research has demonstrated that irrelevant distractors matching the content of memory automatically capture attention (Olivers, Meijer, & Theeuwes, 2006), an effect that can survive (implicitly) learned spatial suppression (van Moorselaar, Theeuwes & Olivers, in press) It should be noted, however, that there was a numerical trend towards reduced distractor-induced activity in the time window of the N2pc in the N2pc and decoding analyses (see Noonan et al., 2016 for a similar finding) Thus, albeit to a lesser extent, distractor repetition may have also affected attentional selection or object individuation. Nonetheless, distractor learning especially modulated distractor-specific processes, as reflected by the repetition-related reduction in the Pd.

Neither target nor distraction location learning modulated early visual stimulus processing, as reflected in the amplitude of the P1 and N1. It is possible that these modulations were masked by overlapping P1/N1 ERPs evoked by the other stimuli in the display. Alternatively, and in line with the longer-latency N2pc en Pd effects, findings from several recent studies also indicate that expectations may modulate only later stages of information processing (Alilović et al., 2018; Rungratsameetaweemana, Itthipuripat, Salazar, & Serences, 2018). Thus, effects of expectation seem to strongly depend on the extent to which the predicted features are relevant for performance, whether positively (as is the case for targets) or negatively (as is the case for distractors). Once the brain has learned that the distractor stimuli can be safely ignored, they no longer need to be inhibited.

Following our ERP findings, target decoding reduced as a function of target repetition in the time window of the N2pc. Note that reduced decoding does not necessarily reflect a change in pattern of neural activity (i.e., location representation), as it could simply be driven by an overall signal attenuation of evoked responses. The correspondence between the latency of our decoding and ERP findings is suggestive of this latter possibility. Distractor decoding did not reveal robust changes as a function of distractor repetition. Yet, we cannot rule out the possibility that with more repetitions, we would have also observed a change in distractor location decoding. Four repetitions may simply not have been insufficient for observing reliable effects.

How does the brain learn that a distractor can be safely ignored? Albeit speculative, rather than modulating activity in regions representing the distractor location, distractor learning may change synaptic efficiency within these regions, analogous to long-term memory and activity-silent coding in working memory (Stokes, 2015). Synaptic memory traces provide a more efficient coding scheme than active suppression through inhibition, and could explain longer lasting effects of learning on how attention is deployed. It would be very inefficient if the brain would need to continually actively suppress responses to irrelevant stimuli that it has learned are safe to ignore. Further research is necessary to test the possibility of distractor learning-related changes in synaptic efficiency.

To conclude, we show that in the context of learning, distractor suppression and target facilitation rely on distinct neural mechanisms. Whereas target learning was associated with increased anticipatory spatial tuning in the alpha-band, distractor learning only reduced reactive inhibition, as reflected in the amplitude of the Pd. These findings argue against direct top-down distractor inhibition, and instead resonate with predictive processing notions in which processing of expected stimuli is suppressed.

## Acknowledgements

This research was supported by a European Research Council (ERC) starting grant (679399) to H.A.S. D.v.M. contributed to design, collected the data, performed the analyses, and contributed most of the writing. HS was closely involved in the design of the experiment and the analysis plan and made significant contributions to the writing. Special thanks to Nathan Coppe for his assistance in EEG data collection.

